# How cultural transmission facilitates a long juvenile learning period

**DOI:** 10.1101/007930

**Authors:** Ryan Baldini

## Abstract

The evolution of the long, slow human life history is a major challenge to evolutionary biologists. A compelling theory states that our late age at maturity allows us to acquire the many skills needed to survive in the economically intensive human foraging niche. Cultural transmission may be a crucial part of this process, by exposing learners to a wealth of information and skills that they would otherwise not likely discover. I use mathematical models to show that whether cultural transmission allows a later age at maturity depends on the details of selection and population regulation. In particular, cultural transmission appears to readily allow a later age at maturity under density-dependent fertility, but may not under density-dependent mortality or density-independent population growth. I close with a discussion of possibilities for future theoretical and empirical research.

## 1 Introduction

The unusually long and slow human life history pattern has provoked much research and debate among evolutionary anthropologists. Even among the primates, humans have an usually long period of juvenile dependency, a late age at sexual maturity, and a long lifespan. Adults reproduce for many years at short interbirth intervals, and, with luck, enjoy a long post-reproductive period (see Kaplan et al., 2000; Mace, 2000; Jones, 2011 for useful reviews of human life history patterns from evolutionary perspectives).

Some researchers have argued that much of this slow life history is the result of selection for an extended juvenile learning period, during which we devote energy to the development of the complex skills needed to survive and reproduce in the extractive human foraging niche (Kaplan et al., 2000). Mastery of extractive foraging skills ultimately reduces adult mortality rates, so that a long juvenile period, long lifespan, and productive adulthood coevolve. This theory is often referred to as the “embodied capital theory” of human life history evolution (e.g. Kaplan et al., 2003). Although I have argued elsewhere that this theory may require further theoretical investigation (Baldini, 2013), the idea is compelling and appears to enjoy some empirical support (Kaplan et al., 2000).

In this paper I argue, similarly to Sterelny (2012), that cultural transmission may be essential to the embodied capital theory, especially with regard to the extended juvenile learning period. Cultural transmission exposes learners to many complex skills and technologies that the vast majority of people could not discover for themselves. These skills require many years of individual practice and experience to develop, but ultimately yield high reward in adulthood. Without culture, there would therefore be little that a young human forager could realistically master, and therefore little reason to extend the juvenile period for so long. I investigate this hypothesis with mathematical models of life history evolution under a variety of selection and population regulation mechanisms. I find that whether culture facilitates a long juvenile period depends on the details of how selection and population regulation operate. Specifically, culture readily facilitates longer juvenile periods under density-dependent regulation of fertility (or infant survival), but may not under density-dependent mortality or exponential population growth. This result suggests that we need to better understand the demographic conditions under which the unique human life history evolved.

The following section develops the argument in more detail and provides an accessible (but not general) mathematical argument for why culture might facilitate a long juvenile learning period. Section 3 contains more explicit models of life history evolution, in which age at maturity and the investment in individual learning coevolve. Because the results of life history models depend strongly on the details of selection and population regulation, section 3 investigates a variety of selective and demographic cases, each of which leads to different qualitative outcomes. The detailed mathematical and numerical results of this section reside in appendices at the end of the paper. Section 4 summarizes the results and suggests directions for future theoretical and empirical research.

## 2 The argument and a heuristic model

### 2.1 The argument

Much of the human maturation process appears to involve the development of complex skills. This is clearly true of modern societies, in which humans spend many years of their lives in formal education systems, even well into reproductive maturity. Subsistence economies, too, require the mastery of many difficult skills. Some evolutionary anthropologists, for example, argue that hunting requires decades of practice to reach peak proficiency, which may not occur until many years after reaching peak physical condition^1^. Individual experience is likely a crucial aspect of this learning process; difficult tasks can’t be learned vicariously. A mathematician can’t master analysis without working through proofs and essentially rediscovering the theorems for himself. Similarly, a forager can’t become proficient simply by observing others; he must log many hours of personal practice before he can support himself and others.

At first glance, the primacy of individual experience seems to discount the role of cultural transmission, because the complex skills needed for survival and reproduction in the human niche can’t be readily transmitted. Yet cultural transmission is essential, because the behavioral domains in which we develop our skills are only available because of the culturally-transmitted information accumulated over generations. Humans spend years learning to use, build, and maintain a complex toolkit and develop complicated skills. Very little of this information is likely to be invented by the young learners themselves.

As an instructive case, consider the bow and arrow. The bow and arrow, with its greater accuracy and stealth than the atlatl, increases an individual hunter’s return rate substantially. Its advantage is so great that its introduction appears to have revolutionized economic and social organization in much of western North America (Bettinger, 2013). The rewards of a bow-and-arrow-based subsistence strategy are likely most enjoyed by those who invest the many hours of practice needed to master the tool’s construction, use, and maintenance. The invention may therefore favor a longer learning period than would simpler technologies, because increased practice continues to pay off. But this long learning period is only possible because hunters are able to culturally inherit the tool and its hunting tactics from previous generations. Of course, the typical human life history was already well established before the invention of the bow and arrow. But the principle should apply to simpler, earlier behaviors. One cannot spend time learning how best to extract honey from a bee hive, use poison to paralyze fish, or weave rain-proof roofing until one is made aware of the possibility to begin with. For all but a few independent inventors, the introduction must occur through social learning. Without such complex tools and techniques, there would be little for a young person to master in the first place, and so little reason to delay reproduction longer than other primates.

This is a somewhat different view of the relationship between social and individual learning than is often presented in the theoretical literature. Usually, theorists assume that social and individual learning are competing learning strategies (e.g. Rendell et al., 2010). While it is true that the information one gathers at any time is either socially derived or not, this view misses the synergy that can arise between social and individual learning. The cultural transmission of complex skills can augment the benefit to individual learning by introducing the learner to otherwise unavailable behavioral domains. My argument echoes Sterelny’s (2012) claim that “human life history characteristics coevolve with technological competence and cultural learning… Extensive cultural transmission supports the development of technological foraging skills that generate extremely rich returns.”

### 2.2 A heuristic mathematical demonstration

Consider the evolution of the age of maturity for some asexual organism. The life course is divided into two learning phases: (1) the learning phase, and (2) the reproductive phase^2^. The learning phase begins at birth, and the reproductive phase begins at some age *a*. For simplicity, assume that the death rate, *µ*, is fixed for all individuals and ages, and that, after maturity, one’s fertility rate, *m*, depends on the amount of information (or individual learning experience), *I*, the individual acquired during the learning phase. *I* is itself a function of *a*; those who spend a longer time learning acquire more information. If we assume further that all strategies are equally strongly regulated by population density through fertility or infant survival, then selection will favor strategies with higher expected lifetime number of offspring, often called *R*_0_ (see appendix A):

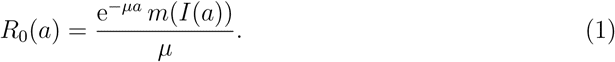

Hence, an increase in *a* requires

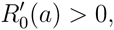

or

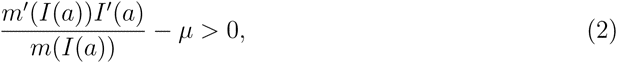

where the prime denotes a derivative. Consider the term *I^′^*(*a*), which is the marginal increase in information gained by extending one’s learning phase beyond the current age *a*. Holding all other things constant, an increase in this derivative causes selection for a longer learning period. My argument is that cultural transmission can increase this derivative by exposing learners to many complex, high-return behaviors and technologies that require years of individual learning to master.

To formalize this argument, suppose that there are *n* possible domains of behavior that one could acquire experience in. These could be conceived as very broad, large-scale behaviors, such as hunting, gathering, fishing, pottery, shelter-building, etc. They could be more narrow, such as “hunting with a bow,” or “hunting species A.” Even narrower, they could represent “processing food with technique B,” “fishing with nets,” or “making tools in style C.” Each of these domains has some probability of being discovered anew by a learner without cultural transmission. Assume for now that this probability of invention is some small number *p*. Assuming (again for simplicity) that inventions occur at a young age, then the number of domains available to a learner *without* cultural transmission on average is simply *np*. Learners then spend their youth developing skill in these domains.

Now consider a population in which learners can observe and imitate the behaviors of others, so that behavioral domains can be learned socially. The probability of acquiring a domain then depends on its frequency in the population, the social learning strategy being used, network size and structure, and so on. I ignore these mechanistic details and simply assume that the probability of acquiring a domain through social learning is an increasing function of the domain’s frequency, *q*, among adults in the population. For example, suppose that the ultimate probability of acquiring some domain is

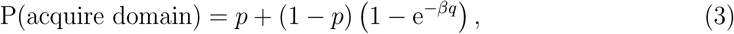

where *β* measures the strength of “cultural selection” (depending on psychological biases, social network composition, and other factors) for the domain. This equation says that, in a cultural population, one can learn the behavior either through individual invention (with probability *p*), or, if not (1 − *p*), then through social learning (1 − exp(−*βq*)). Because *q* is itself an evolving variable, this probability changes through time as more or fewer people become aware of the behavioral domain. This will tend to settle down on some equilibrium value 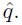 Although equation (3) can’t be solved for equilibrium in elementary terms, it is clear that 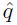 will be larger than *p* because of the additional probability of learning behavioral domains socially. It may be much larger if social learning is effective and the domain is very difficult to discover individually. Because there are more known skills to develop, the returns to individual learning diminish more slowly that in an acultural population, and *I^′^*(*a*) will likely be greater.

It is possible that the shift toward social learning reduces individual inventiveness such that *p* will be reduced to some value *p^*^ < p*. In that case, one can show that 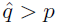 only if

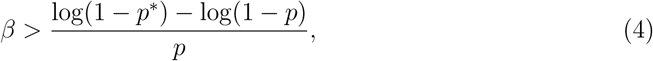

which, if *p*, *p^*^*, and their difference are all small, can be reasonably approximated by

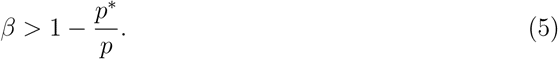

The condition *β >* 1 is required for a social learning population to maintain a domain in the population in the first place, so the above inequality is reasonable.

For the remainder of the paper, I do not treat the evolution of cultural transmission itself. Rather, I focus on how an increase in 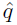 affects life history evolution. The implication is that cultural transmission will tend to increase 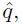 based on the above argument.

## 3 Results of complete models

A more complete analysis should track not only the genetic evolution of the age at maturity, but also the amount of energy invested in individual learning itself, as this is likely to change as the learning period changes. In other words, everything else is *not* held constant in inequality (2). Let the variable *α* represent the amount of energy or effort devoted to individual learning. An increase in *α* increases the rate at which information is learned, but comes at a cost to either survival or fertility (see cases below) because of the energy needs of maintaining the cognitive structures that allow for learning, or because the time-consuming act of learning itself may reduce survival or fertility.

For the individual learning function *I*, I assume a diminishing-returns function:

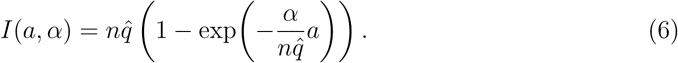

The derivation is as follows: an individual devotes an amount of energy *α* to individual learning. This is divided evenly into all the domains available to the learner - of which there are 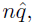 on average. The rate of learning diminishes exponentially toward an asymptote of 1 for each domain. When added across all domains, this implies a maximum total *I* of 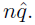 Learners never quite reach this maximum value; they may fall well short of it if the length of their learning period is short. Figure 1 shows the general shape of *I*(*a*) for different values of 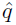 and *α*.

**Figure 1.**
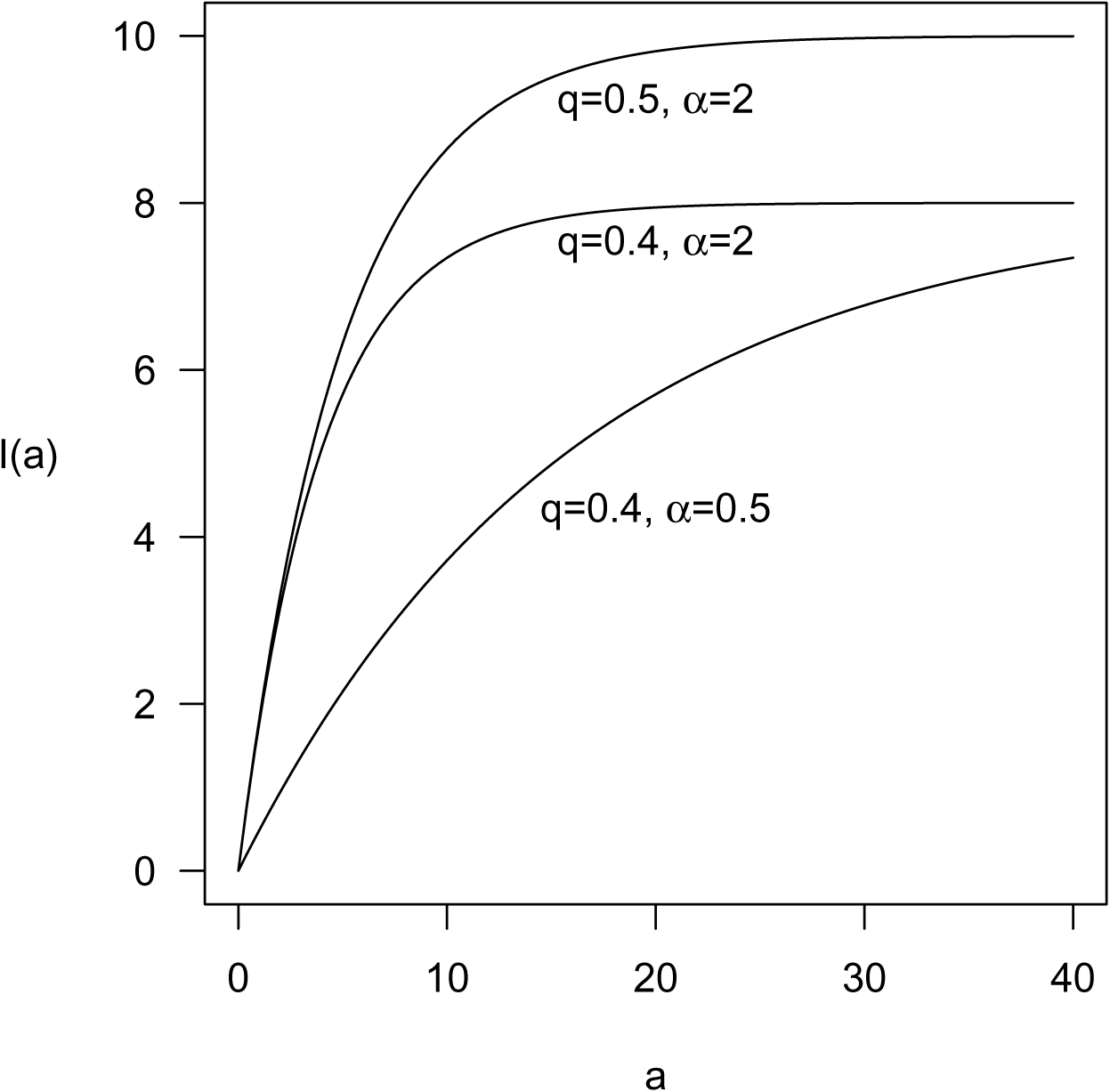
Learned information *I* as a function of length of learning stage *a*. Larger *α* allows faster learning; larger *q* allows more to learn. *n* = 10.

**Figure 2:**
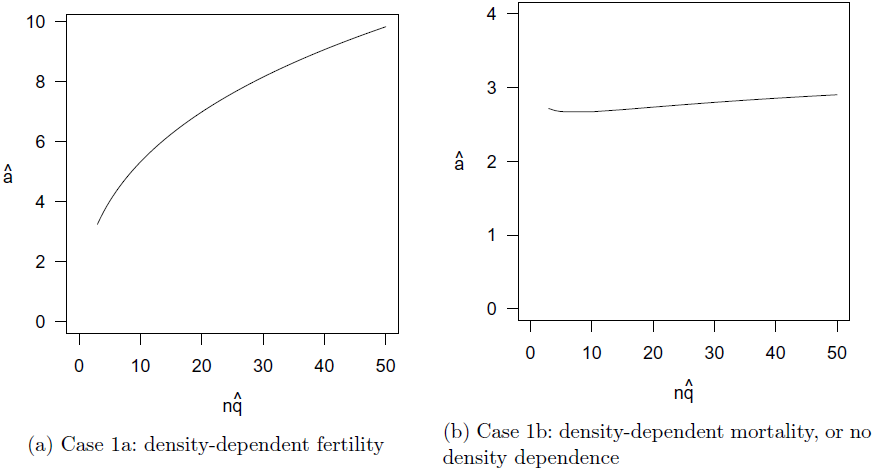
Equilibrium age at maturity (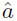) under various numbers of available learning domains (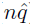) when selection acts on fertility. *µ* = 1*/*30, *c* = 1*/*30, *s* = 0.02. The unplotted range for small 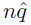 corresponds to population extinction.

I consider simultaneous evolution in *a*, the age at maturity, and *α*, the energy devoted to individual learning. In general, the action of selection depends on how vital rates (mortality or fertility) are affected by learning, and how the population grows and is ultimately regulated (Charlesworth, 1994). The heuristic argument above only considered selection and density dependence acting on fertility. Below I allow two simple kinds of selection: case 1, where increased information leads to higher fertility, and case 2, where increased information leads to lower mortality. Within these cases I allow three kinds of population regulation: density-dependent fertility, density-dependent mortality, and unregulated, exponential population growth. Under the conditions explained in Appendix 1, the latter two forms of growth always select for the same strategy, so I combine them in the following analysis. In reality, density may affect *both* mortality and fertility. In that case, the evolutionary equilibrium lies somewhere between the two extremes (Baldini, 2013).

### 3.1 Case 1: selection on fertility

First consider the case where information gained through learning increases one’s fertility rate, but not one’s mortality rate. That is, skills acquired in learning increase an adult’s ability to produce surviving offspring - perhaps by better acquiring resources, attracting mates, or providing proper care for infants. I assume a simple linear effect of information *I* on the fertility rate, *m*, so that equation (6) becomes

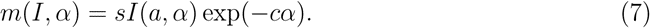

This assumes that an individual with no learned information at all has zero fertility.

#### Case 1a: density-dependent fertility

Suppose that the fertility rates of all strategies decrease as population density grows. The appendix contains the mathematical details of this model, and derivations of the selection gradients on *a* and *α*. The general result arises that there is a single positive optimum, denoted by 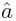 and 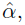 and any increase in 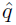 *always* selects for larger 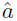 and 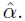 Thus, if cultural transmission exposes people to more behavioral domains, selection will favor longer, more intensive juvenile learning periods under Case 1a conditions.

Figure 3a shows the typical shape of 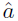 as affected by 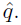 Since 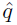 is always multiplied by *n* in the model, figure 3a plots 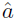 on this product, which can be interpreted as the number of behavioral domains in which the average learner can gain experience. The effect of 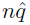 can be large, especially compared to the case of density-dependent mortality or exponential growth (figure 3b; see just below).

**Figure 3:**
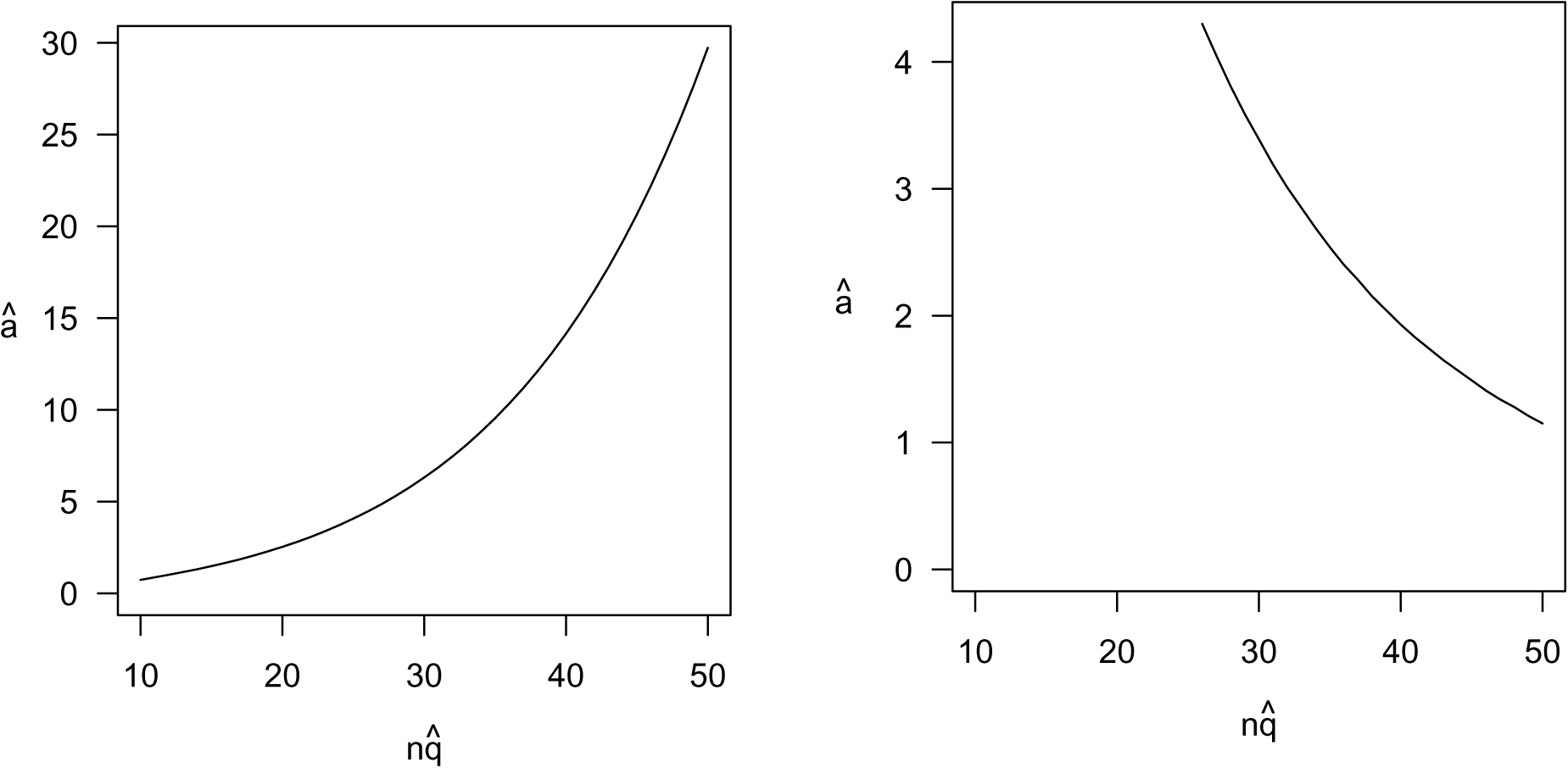
Equilibrium age at maturity (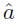) under various numbers of available learning domains (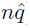) when selection acts on mortality. *µ*_0_ = 1, *c* = 1*/*30, *s* = 0.1, *m* = 1. The unplotted range corresponds to population extinction.

#### Case 1b: density-dependent mortality, or exponential growth

Now suppose that either the mortality rate increases linearly with population density, or neither mortality nor fertility are density-dependent. These two cases select for the same life history schedule under the simple functional forms shown in Appendix A. Unfortunately, I have been unable to derive any general result about the effect of 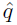 on 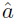 or 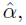 as the gradients are more complicated. For any particular set of parameters, however, we can solve for equilibrium numerically. Appendix B contains tables for many such equilibria under various parameter values.

In every case analyzed, there was only one positive equilibrium 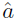 and 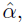 and the effect of 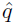 on 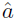 was about the same. For small 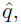 an increase in 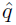 causes 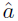 to decrease. At some critical point in 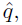 the effect reverses, so that further growth in 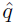 causes 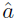 to grow as well. Figure 3b shows an example of this effect. Thus, the effect of behavioral domain number on optimal age at maturity appears to be equivocal. Furthermore, the effect is generally weaker in case 1b than in all other cases. I have not been able to explain on intuitive grounds why the differences arise.

### 3.2 Case 2: selection on mortality

Now consider the case where information gained through learning decreases one’s mortality - or, in other words, lengthens one’s lifespan. A difficulty arises here, because the learner’s mortality rate should then change over time as the learner acquires more information. I ignore this complication by simply assuming that an individual has a constant death rate determined by the amount of information that he will ultimately acquire. One justification of this is to imagine that one’s juvenile birth rate is a function of his parent’s skill level, which, because of asexual reproduction, is exactly that which the juvenile will eventually acquire. This justification is admittedly contrived, and the functional form I choose probably strongly affects the results of selection. I leave more detailed investigation of mortality patterns to future research.

The mortality rate must be positive, so I assume a negative exponential effect of *I* on *µ*, and a positive linear cost to individual learning:

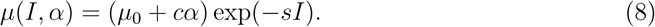

*µ*_0_ is the baseline mortality rate for a learner with *I* = *α* = 0. I simply let *µ*_0_ = 1, so that the majority of individuals born into a lineage with 0 knowledge would die within 1 year of birth. *I* is again given by equation (7). Note now that *m*, the fertility rate, is fixed for each strategy.

Unlike Case 1, Case 2 always has a stable equilibrium at 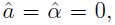 where individuals forgo a juvenile learning period entirely (see appendix A). Instead, they mature as soon as they are born (or, more realistically, as early as is physiologically possible, considering other constraints on the organism). There is usually also a positive stable equilibrium, with a juvenile learning period, but its existence is not guaranteed. This means that the long-term result of selection may depend on the population’s initial state. In particular, selection on mortality alone cannot get a short-lived population “off the ground,” but, once past a critical threshold, the population may evolve a longer juvenile period. I consider this 0 equilibrium to be unrealistic for humans and primates: clearly humans have long juvenile period. Thus I focus below on the positive equilibrium, when it exists.

#### Case 2a: density-dependent fertility

Density dependence operates as in Case 1a. Appendix A contains the selection gradients on *a* and *α*. I could not derive general results about 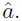 Nonetheless, the tables in Appendix B consistently show that increasing 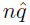 selects for a longer juvenile period 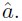 The effect can be very strong, and always has the same general shape. An example is shown in figure 3a. The strength of the effect notably increases as the effect of information on mortality increases (i.e. as *s* increases). In other words, the more one can lengthen one’s lifespan by learning, the later is the optimal age at maturity.

#### Case 2b: density-dependent mortality, or exponential growth

Appendix A contains the selection gradients. Again, I could not derive general results, but the tables in Appendix B suggest that increasing 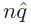 always selects for a *shorter* juvenile period 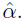 This is opposite to Case 1a and 2a. The effect appears to be strong; figure 3b shows an example.

## 4 Summary, problems, and possible extensions

Table 1 summarizes the basic qualitative results of my analysis.

**Table 1:** The effect of increasing 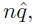 the number of behavioral domains available to learners, on 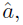 the optimal age at maturity. Arrows indicate the directional effect. Each cell indicates whether the effect is relatively strong or weak. Only the top left cell is proven.

|  |  | Density Dependence: |  |
| --- | --- | --- | --- |
|  |  | Fertility | Mortality |
| Selection: | Fertility | $\hat{a} \uparrow$<br>strong | $\hat{a} \uparrow$ or ↓<br>weak |
| | Mortality | $\hat{a} \uparrow$<br>strong | $\hat{a} \downarrow$<br>strong |

One conclusion is certain: the effect of cultural transmission on the evolution of age at maturity depends on the nature of population regulation. Sometimes the effect is in the predicted direction, with cultural transmission facilitating later maturity, and sometimes not. I found a similar dependence on demographic conditions when reanalyzing an earlier model of embodied capital theory (Kaplan et al., 2000; Baldini, 2013). Together, these papers prove that our understanding of human life history evolution cannot proceed much without a better understanding of the demographic conditions faced by our ancestors.

A tenuous pattern found here is that selection readily favors later maturity under density-dependent fertility, but not under density-dependent mortality or density-independence. A possible explanation of this is as follows: The age at maturity is strongly affected by lifespan, because dying before maturity implies zero fitness. Under density-dependent evolution, selection always leads to a larger equilibrium population size, which in turn leads to higher mortality and a shorter lifespan. This effect may cancel out any positive effect of selection on investment in lifespan. The corresponding effect for exponentially growing populations is this: as selection favors strategies that grow ever faster, the reproductive values of older ages become ever more discounted, weakening the strength of selection for delayed reproduction. This effect does not arise under density-dependent fertility, because there the mortality rate either stays the same or shrinks as the population acquires more information and grows.

I caution, however, against taking this pattern too seriously. More theoretical work is needed. I have only considered the simplest functional forms for vital rates and density-dependence - some of which are unrealistic for primates. The assumptions of a constant mortality rate and a constant fertility rate following maturity do not fit primate life history patterns. Furthermore, I assumed that density-dependent mortality affects all ages equally, but this is probably not true (see, e.g., Wood and Smouse, 1982). I did not consider variation in the effect of population density on different strategies, which essentially rules out the possibility of any *r* vs. *K* selection effects. On a more technical level, I have not demonstrated that each parameter combination investigated actually causes a stable demographic equilibrium - an important concern for the density-dependent models in particular, as cycles and chaos may arise. Nor could I investigate evolutionary stability rigorously, as this requires explicit genetic models.

A possibly important omission of this paper is to not allow cultural transmission itself to evolve. Instead, it acts as an exogenous parameter through 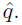 A simple way to resolve this would be to let *β* (of equation (3)) evolve similarly to *α*, by imposing a cost to higher values. Large *β* would imply more social learning. I ignored this detail here for two reasons. First, much has been written about the evolution of cultural transmission itself (see reviews in Henrich and McElreath, 2003; Richerson and Boyd, 2005; Boyd et al., 2011), though I know of none that considers its effect on life history. Second, because culture transmission does not directly interfere with individual learning in the present theory, there is little reason to believe that it wouldn’t evolve in the first place. Still, it is possible that including social learning in the organism’s energy budget would alter the results; future work should check this.

Altogether, the most valuable next step for theoretical research might be to use more primate-or human-typical life history schedules in our theoretical models. The demographic conditions under which the modern human life history evolved must lie somewhere in the range spanned by modern human populations and our primate relatives. Investigating this range will constrain our attention to only the most relevant conditions, and has already revealed some possible generalities about human evolution (Jones, 2009). In the empirical domain, we desperately need a better quantitative understanding of how human populations are regulated in subsistence economies. Little direct research on humans exists. I know only of the empirical study by Wood and Smouse (1982) on a New Guinea population, which suggested that density-dependent mortality may act strongly on the youngest and oldest age classes. The large literature on population regulation among other large mammals (see a review in Gaillard et al., 1998) may provide useful methods and hypotheses.

1 See Kaplan et al. (2000); Walker et al. (2002); Walker and Hill (2003); Gurven et al. (2006). There is a debate, however, whether patterns in human foraging productivity across age reflect skill acquisition or simply physical growth and senescence. For an argument supporting the latter, see Bliege Bird and Bird (2002); Bird and Bliege Bird (2002); Blurton Jones and Marlowe (2002).

2 Of course, humans can continue to learn after they reproduce. Ideally one would want to represent the effort contributed to learning and to reproduction with real numbers, rather than dividing it into discrete stages. I do the latter here for simplicity; future investigations should check that the same basic results arrive. Kaplan et al. (2000) do include a learning process after reproduction, but simply assume that it is a an unevolving process of exponential skill growth, which seems unrealistic.

## A Selection gradients for *a* and *α*

### Case 1a

Suppose that the total population density, *N*, reduces the fertility function, *m*, via an exponential multiplier:

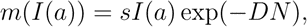

where *D* measure its affect on fertility. Selection favors the strategy in *a* and *α* which produces the largest carrying capacity (Charlesworth), subject to the constraint

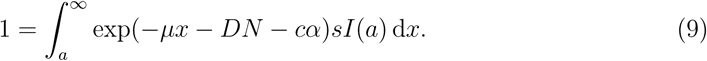

The term exp(−*DN)* can be removed from the integral, so maximizing *N* is equivalent to maximizing the expected lifetime number of offspring at *N* = 0, often called *R*_0_:

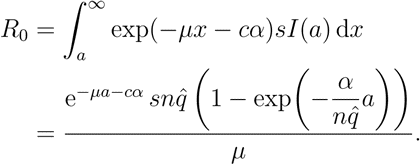

The second line follows from the form of *I* assumed in the text.

The selection gradients for *a* and *α* are the derivatives of *R*_0_, which are proportional to

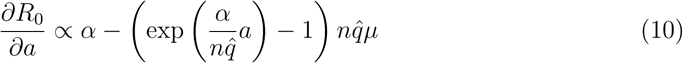

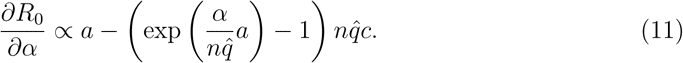

The equilibrium values, 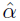 and 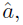 are found by setting these equal to zero and solving for the two variables. That is, they solve the equations

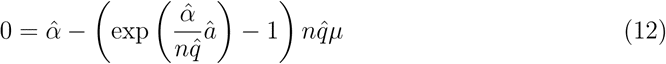

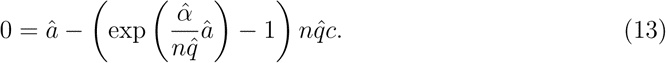

These cannot be solved in elementary terms, but we can prove in general that 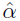 and 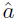 always increase with 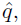 and therefore will be larger in a social learning population.

To see this, multiply the expression in (11) by 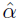 and the expression in (12) by 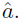 The resulting expressions are still zero, so we may set them equal to each other:

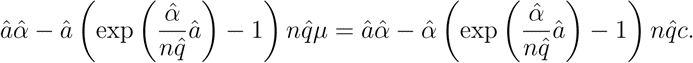

Simplifying this shows that

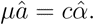

Plugging this result into (11) gives us

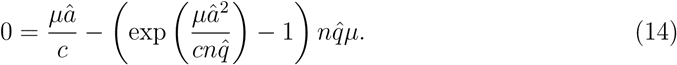

This equation has a unique positive solution in 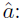 the quantity on the right has a root at 0, increases from 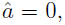 has only one critical point, and approaches 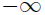 as 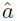 approach 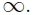 This means that the right hand side of (14) reaches a single positive maximum after which it decreases indefinitely, crossing zero at some positive value for 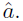 Its derivative with respect to 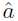 must be negative at the positive root - a fact which is used below.

Expressing 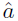 as a function of *q*, implicitly differentiating (13) for *q*, and solving for 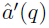 gives us

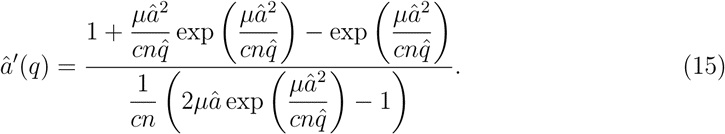

I prove that this is positive by showing that the numerator and the denominator are both positive. First, the numerator of (14) is an expression of the form

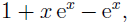

where *x*, in this case, is positive. Expressing e*^x^* as a Taylor series allows the following manipulations:

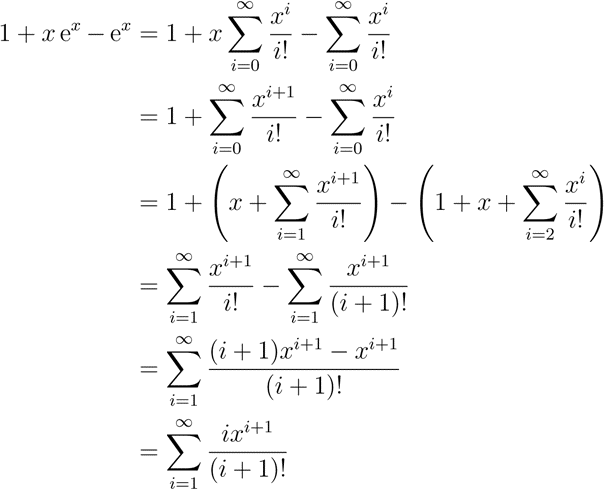

Because *i* and *x* are always positive, the sum is always positive. Second, the denominator of (15) is proportional to the negative of the derivative of the right hand side of (14) with respect to 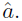 I proved above that the derivative is negative at 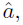 so the denominator of (15) must be positive. Therefore 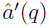 is positive and, because, 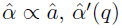 is also positive.

The trivial equilibrium 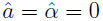 is unstable and, in fact, impossible, because it implies zero fertility and therefore extinction. For small *a* and *α*, *R*_0_ *≈ aαs/µ*. Selection always favors larger *R*_0_, so *α* and *a* always evolve upward from very small values.

### Case 1b

Suppose that the mortality rate increases linearly with density, again by an amount *D*. Then selection maximizes *N*, subject to the equation:

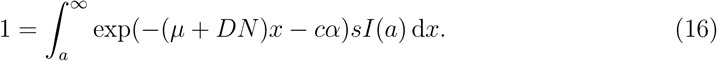

In this case, the term involving *N* cannot be removed from the integral, so selection does not favor the same strategy as in case 1a; we cannot simply maximize *R*_0_. We must instead maximize *N* above.

Alternatively, consider the case without density-dependence, such that a population approaches a constant growth rate *r*. Then the Euler-Lotka equation is

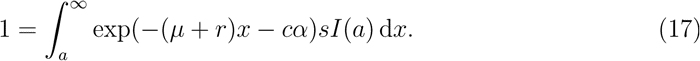

A strategy that grows exponentially faster than all others will eventually approach fixation, so selection maximizes *r*. But notice that *r* of equation (16) has the same functional role as *DN* in equation (15). Thus, selection favors the same life history strategies in both cases. I use the density-independent notation (*r*) here, but the following results apply to both.

The solution to (16) in *r* is

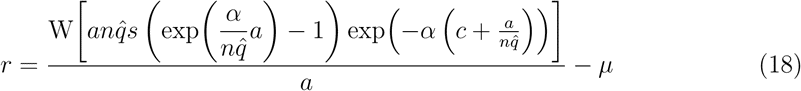

The W[*] term denotes a product-log, defined implicitly by the equation *x* = W[*x*] e^W[^*^x^*^]^. It is positive whenever its argument is positive, which is the case here. The selection gradients for *a* and *α* are then proportional to

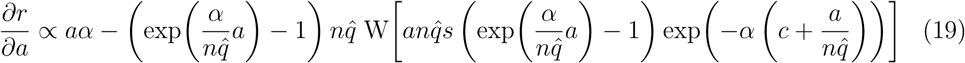

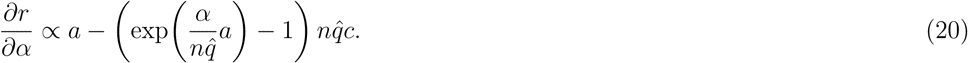

As in Case 1a, the equilibrium 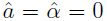 is unstable and causes population extinction. For small *a* and *α*, *r ≈ aαs − µ*, so *r* increases with *a* and *α* when they are small, implying positive selection.

### Case 2a

By the same logic as in Case 1a, but now applied to mortality selection, we have

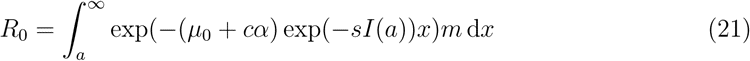

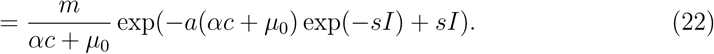

The gradients in *a* and *α*, using *I* from equation (7), are then

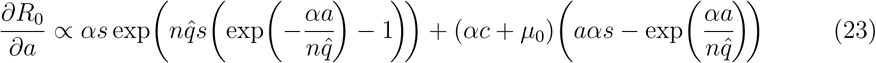

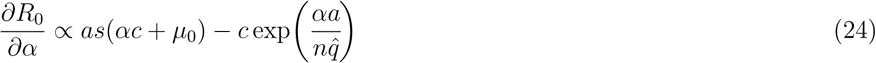

A population at equilibrium 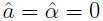 avoids extinction if *m > µ*_0_. It is stable whenever it exists: the gradients for *a* and *α* are then −*µ*_0_ and −*c*, respectively. Searching for roots numerically shows that there are often two positive equilibria, of which only one can be stable. The paper focuses on this stable positive equilibrium.

### Case 2b

By the same logic as in Case 1b, but now applied to mortality selection, we have

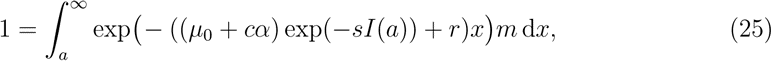

The solution in *r* is

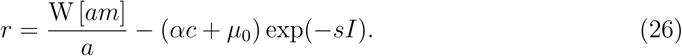

Using *I* from equation (7), the gradients in *a* and *α* are

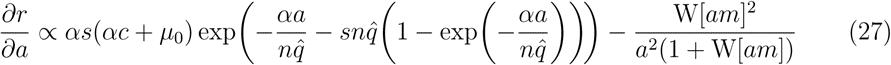

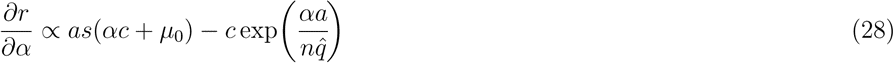

A population at equilibrium 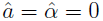 exists and is stable under the same conditions as Case 2a. There are, again, often two positive equilibria, only one of which may be stable.

## B Tables in 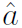

### Fertility selection

The following tables show the equilibrium age at maturity (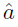) for cases 1 and 2. Each table shows how variation in 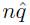 and some other parameter affect 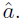 Variables not shown in each table are kept at a baseline value, as follows: *s* = 0.05, *µ* = 1*/*30, *c* = 1*/*30. Empty cells indicate that the population goes extinct under that set of parameter values.

#### Case 1a

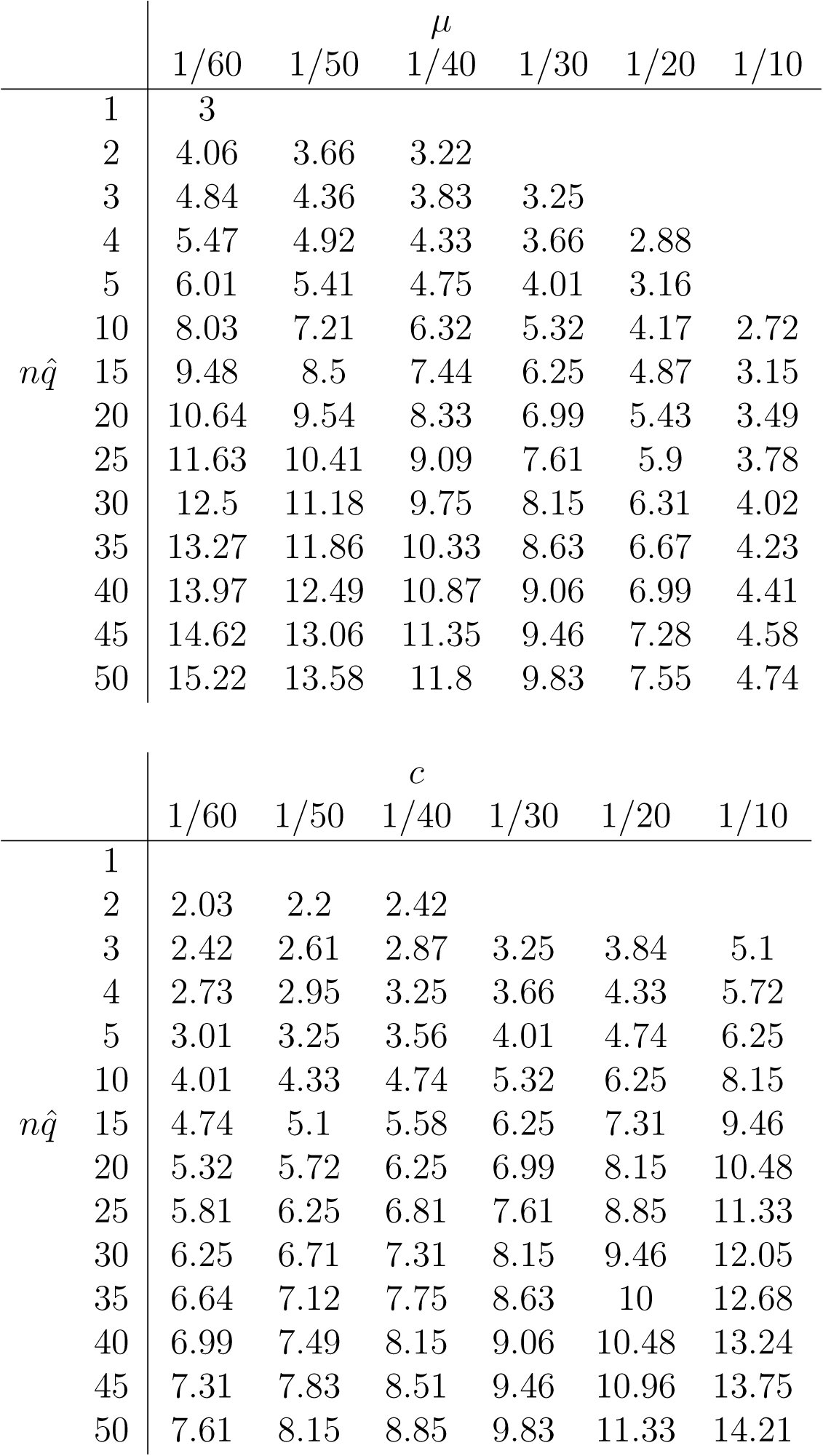

#### Case 1b

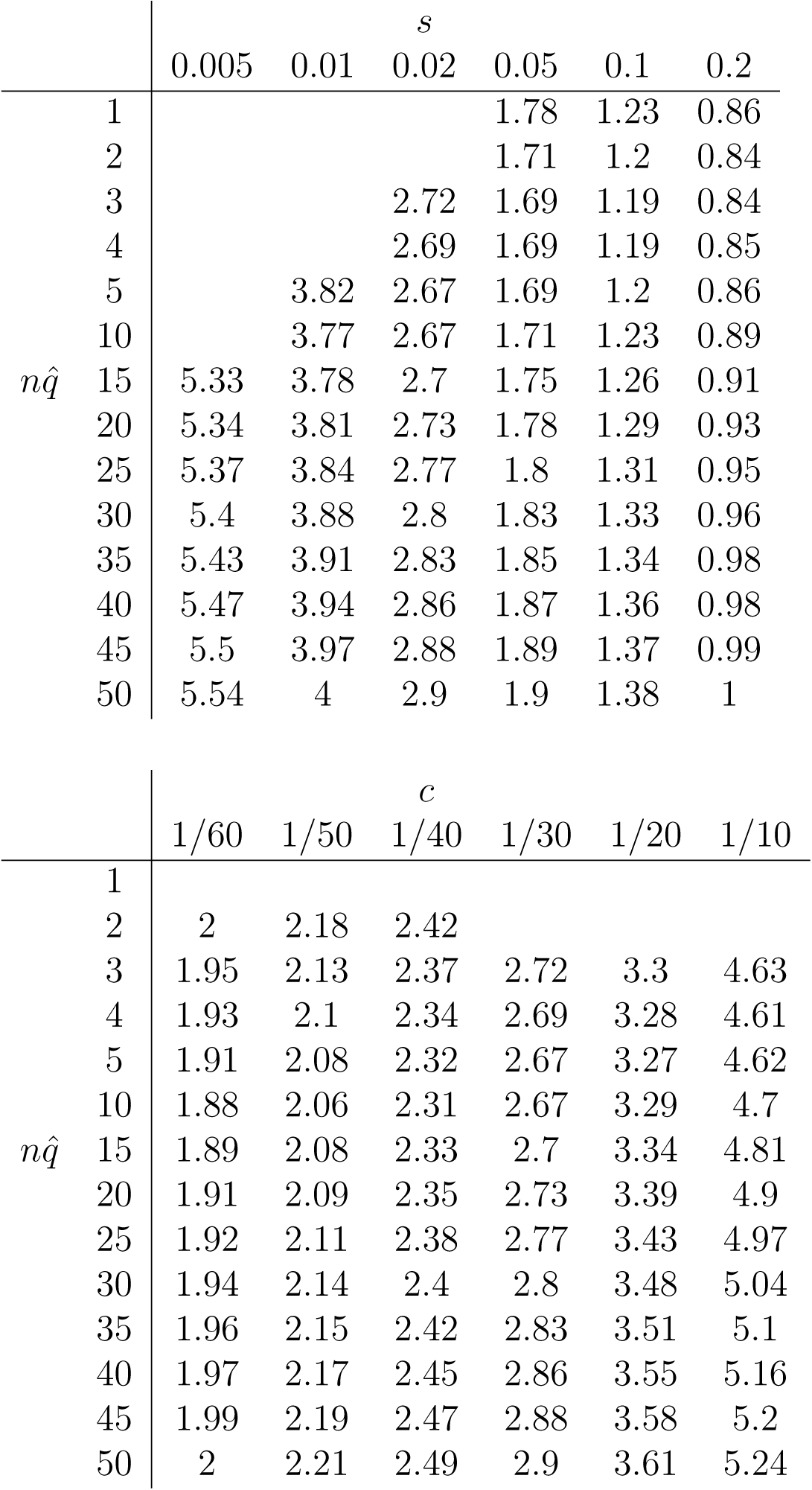

### Mortality selection

#### Case 2a

0 implies that there is no positive evolutionary equilibrium. *µ*_0_ = 1, *c* = 1*/*30, *s* = 0.1.

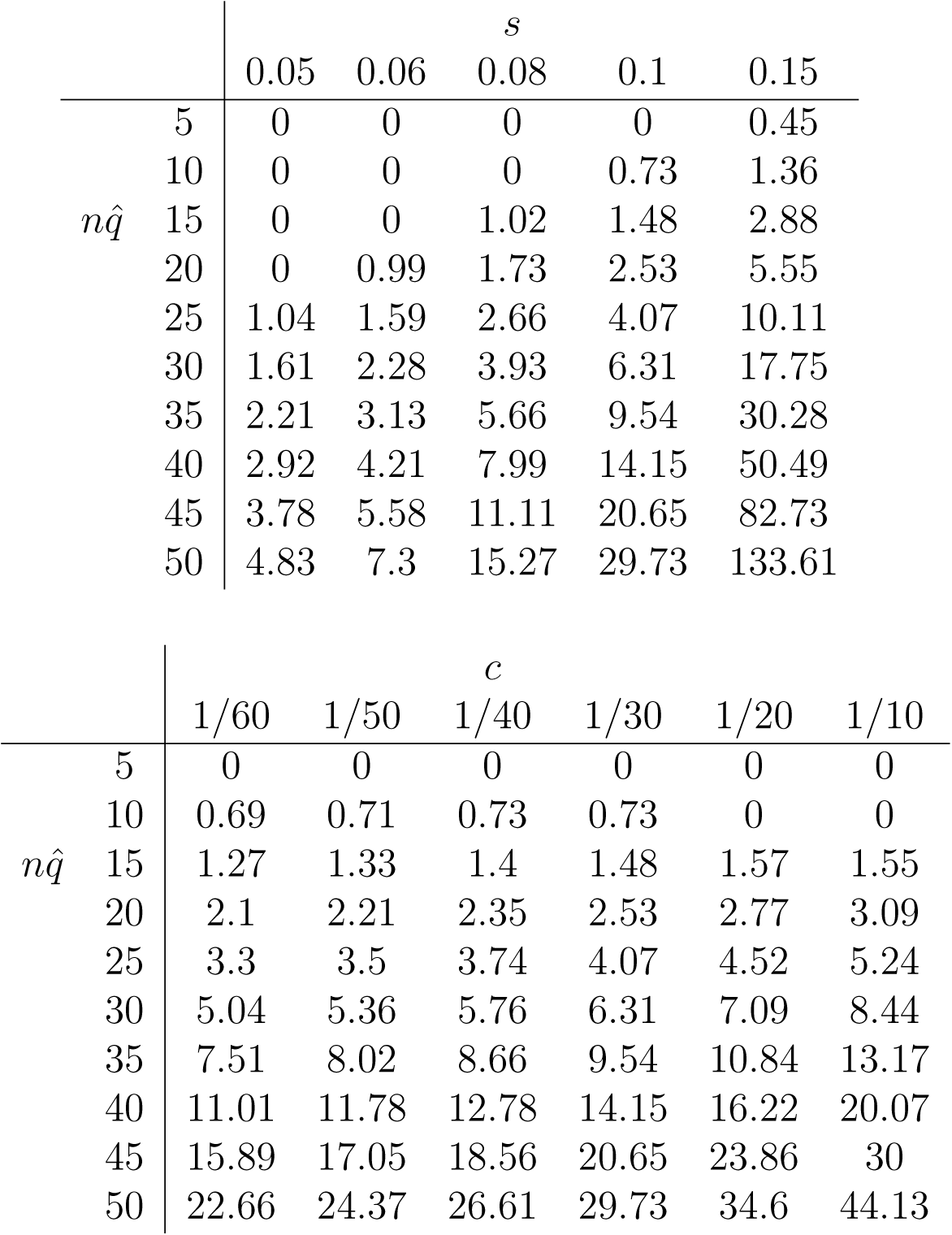

#### Case 2b

Baselines: *m* = 1*/*4, *µ*_0_ = 1, *s* = 0.1. Blank values indicate that the population goes extinct for that parameter combination.

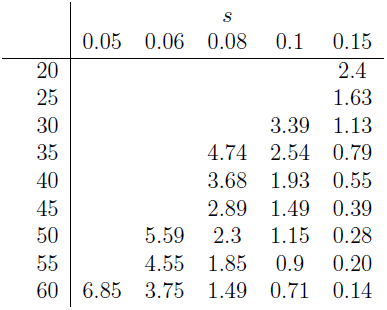

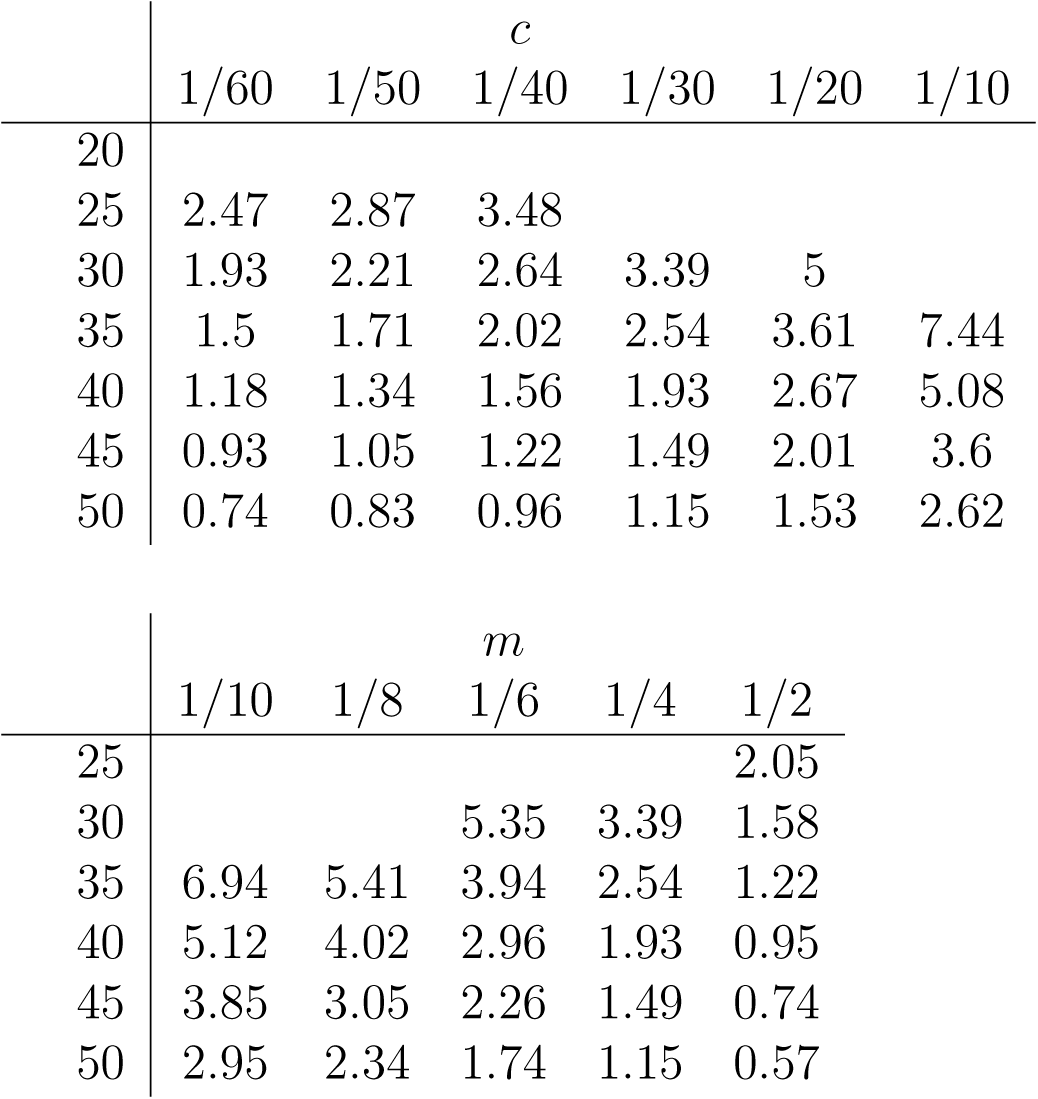

